# Alternative Promoters Drive Human Cytomegalovirus Reactivation from Latency

**DOI:** 10.1101/484436

**Authors:** Donna Collins-McMillen, Mike Rak, Jason Buehler, Suzu Igarashi-Hayes, Jeremy Kamil, Nat Moorman, Felicia Goodrum

## Abstract

Reactivation from latency requires reinitiation of viral gene expression and culminates in the production of infectious progeny. The major immediate early promoter (MIEP) of human cytomegalovirus (HCMV) drives the expression of crucial lytic cycle transactivators but is silenced during latency in hematopoietic progenitor cells (HPCs). Because the MIEP is poorly active in HPCs, it is unclear how viral transactivators are expressed during reactivation. We demonstrate that transcripts originating from alternative promoters within the canonical major immediate early locus are abundantly expressed upon reactivation, whereas MIEP-derived transcripts remain undetectable. Further, we show that these promoters are necessary for efficient reactivation in primary CD34+ HPCs. Our findings change the paradigm for HCMV reactivation by demonstrating that promoter switching governs reactivation from viral latency in a context-specific manner.

---

Reactivation of latent human cytomegalovirus (HCMV) infection poses a life threatening risk to immunocompromised individuals, such as stem cell or organ transplant recipients and those that become immunosuppressed during treatment for cancer (1). While the HCMV replicative cycle has been studied extensively, our understanding of the mechanisms controlling the entry into and exit from latency is far from complete.

During productive infection, the HCMV genome is transcribed in a temporal cascade comprised of three kinetic classes of gene expression designated as immediate early (IE), early (E), and late (L) (2). The IE proteins, in particular IE1 and IE2, play critical roles in initiating the HCMV lytic cycle by transactivating the expression of cellular and viral genes, and suppressing the innate immune response (3). During latency, IE1 and IE2 expression is silenced, resulting in diminished viral gene expression and the absence of productive replication (4). Signals that stimulate reactivation induce IE1 and IE2 expression to allow re-entry into the viral replicative cycle (5). Regulation of IE1 and IE2 expression is thus a pivotal event controlling the switch between latent and reactivated states.

The MIEP is highly active in cells permissive for lytic HCMV replication, during which it drives high level expression of mRNAs encoding IE1 and IE2 (6–9). In cell types that support HCMV latency, however, such as CD34+ human progenitor cells (HPCs) and CD14+ monocytes, the MIEP is silent (10–12). Because signaling events that trigger HCMV reactivation also increase IE1 and IE2 mRNA levels (13–17), it has been presumed that de-repression of the MIEP is critical to reinitiate the viral lytic cycle. However, the origin of IE1 and IE2 transcripts during HCMV reactivation has not been formally defined.

To determine if the MIEP does in fact drive the accumulation of IE1 and IE2 mRNAs during reactivation from latency, we measured the expression of IE1 and IE2 transcripts during experimental latency and reactivation in the monocytic THP-1 cell line. THP-1 cells are an established model for studying HCMV latency and reactivation (18–20) in which reactivation can be synchronized. Cells were infected with HCMV (TB40/E strain) and allowed to establish latency for 5 days post infection (dpi). To induce reactivation, latently infected cells were treated with 12-O-tetradecanoylphorbol-13-acetate (TPA), which promotes monocyte-to-macrophage differentiation and triggers viral reactivation. The IE1 and IE2 proteins were detected immediately after infection but decreased from 2 to 5 dpi to levels below the limit of detection (Figure 1A), consistent with the establishment of latency. IE1 and IE2 proteins accumulated rapidly following TPA treatment, consistent with viral reactivation. Parallel cultures treated with the solvent control (DMSO) maintained latency, and levels of IE1 and IE2 proteins remained below the limit of detection. The accumulation of IE1 and IE2 transcripts following infection and reactivation in THP-1 cells was measured using quantitative reverse transcriptase PCR (RT-qPCR) and primers detecting spliced IE1 and IE2 transcripts (Figure 1B, black arrows). The expression of IE1 and IE2 transcripts corresponded to proteins detected at 1 dpi and following reactivation (Figure 1C). Surprisingly, using a primer pair to monitor transcription from the MIEP or the 5’ distal promoter (Figure 1B, orange arrows), we detected few transcripts derived from the MIEP/dP at any time after infection or reactivation (Figure 1C), although MIEP-derived transcripts accumulated robustly during productive infection of fibroblasts (Figure S1). The absence of MIEP-derived transcripts following reactivation in THP-1 cells indicates that the re-expression of IE1 and IE2 during reactivation is driven by alternative promoters.

**Fig 1.**
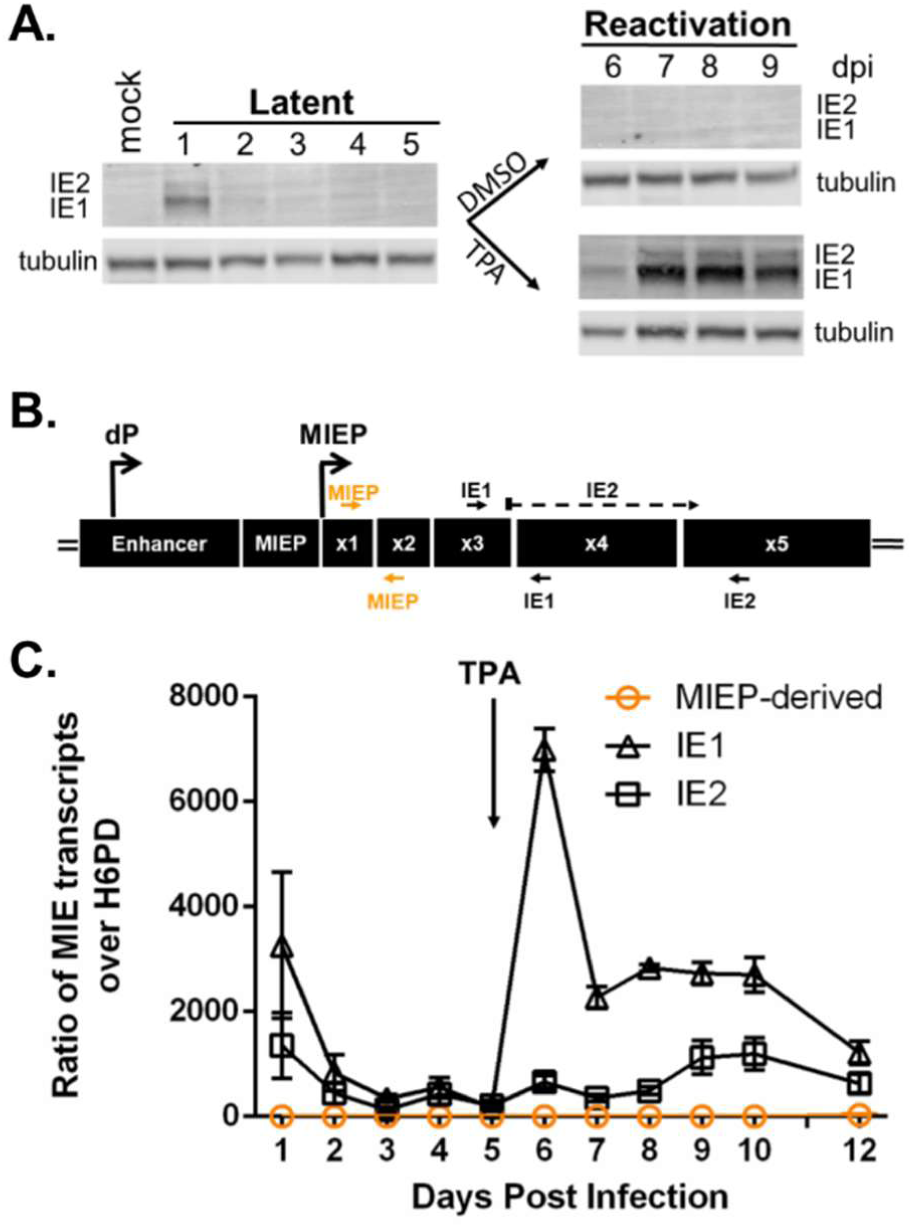
IE1 and IE2 are expressed during HCMV reactivation but do not arise from the MIEP. **A)** THP-1 acute monocytic leukemia cells were infected with TB40/E WT HCMV (MOI = 2) and cultured for 5 days to allow the establishment of latency. At day 5, cells were treated with TPA (to promote macrophage differentiation and viral reactivation) or with a DMSO control. Whole cell lysates were collected at the indicated timepoints and IE1 and IE2 protein accumulation was measured by immunoblotting using a monoclonal antibody recognizing both IE1 and IE2 (kind gift, Tom Shenk). Tubulin was used as a loading control. A single experiment (representative of three independent experiments) is shown. **B)** Schematic of the major immediate early (MIE) locus. The distal promoter (dP) and major immediate early promoter (MIEP) and primers to detect IE1 (spanning exons 3-4), IE2 (spanning exons 3-5) or MIEP/dP-derived transcripts (including exon 1-2, orange) are indicated. C) IE1, IE2 and MIEP/dP-derived transcripts were detected over a time course following infection and reactivation in THP-1 cells by RT-qPCR. Transcripts are quantified as a ratio over the low copy housekeeping gene hexose-6-phosphate dehydrogenase (H6PD). Data from three independent replicates is shown; error bars indicate standard deviation.

We recently identified two promoters within an intron 3’ of the MIEP (intronic promoters, iP1 and iP2) that drive the expression of alternate transcripts encoding IE1 and IE2 (Figures 2A, 2B) (21). iP-derived transcripts differ from MIEP-derived transcripts in that they lack the non-coding exon 1 but encode the complete set of distal exons necessary to express full length IE1 or IE2 proteins. Moreover, the intronic promoters are dispensable for viral replication in fibroblasts (21). Given the minimal activity of the MIEP in hematopoietic cells, we hypothesized that the iP1 and iP2 might contribute to IE1 and IE2 expression during reactivation. We therefore monitored the accumulation of iP1 and iP2-derived transcripts in THP-1 cells using primer pairs that discriminate between the distinct MIE transcripts (Figure 2B; sequences in Table S1). We found that MIE transcripts were most abundantly derived from iP2 between 1 and 5 dpi and were robustly induced from iP2 following reactivation (Figures 3C, S2). iP1-derived transcripts also increased after reactivation, but to a lower level. Although a small number of MIEP-derived transcripts were detected at 10 – 14 days following reactivation (Figure S2), the MIEP appeared to be far less active than either of the intronic promoters. Thus, transcripts originating from iP1 and iP2 account for the overwhelming majority of IE1 and IE2 transcripts in THP-1 cells, suggesting that they play critical roles in the re-expression of IE proteins (Figure 1A) and reactivation from latency in hematopoietic cells.

**Fig 2.**
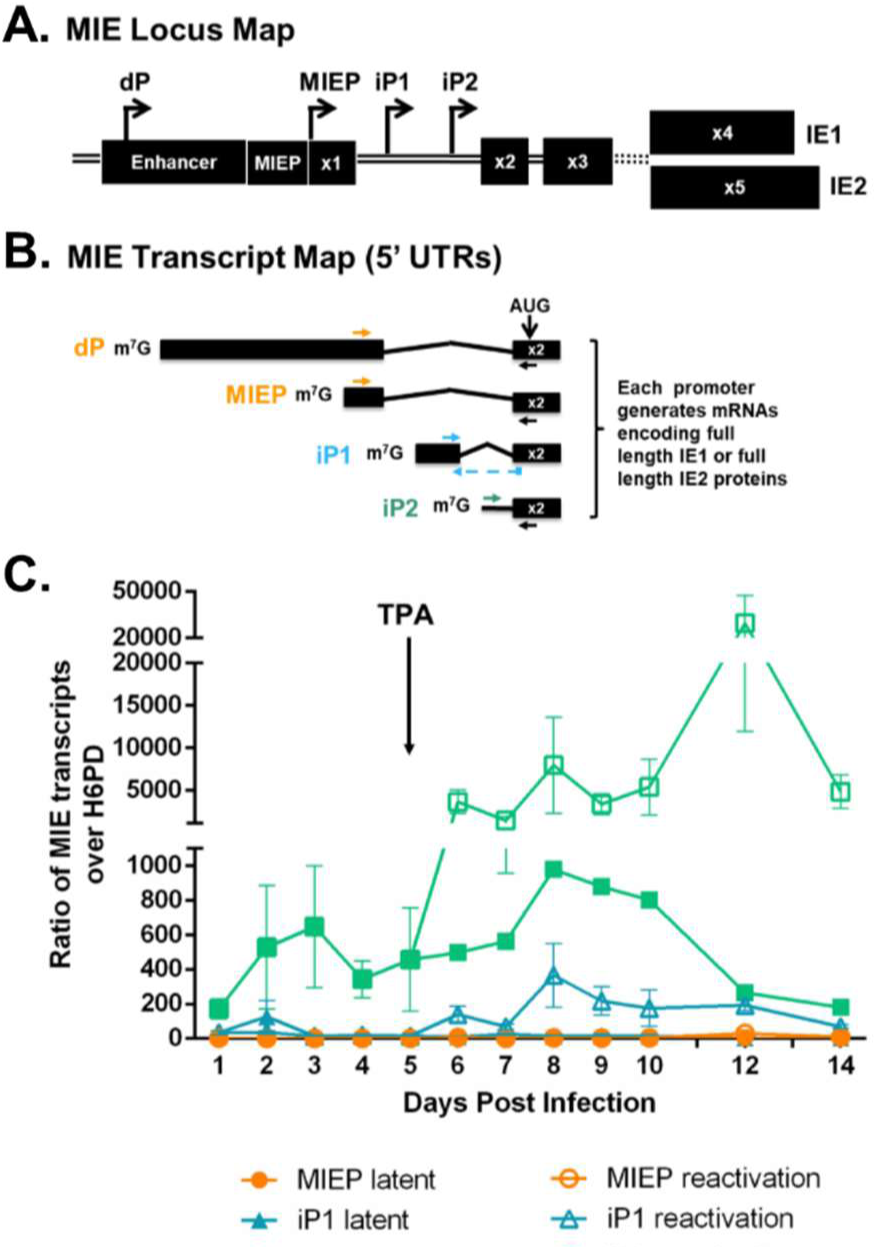
IE1 and IE2 transcripts derived from intronic promoters predominate during reactivation. **A)** Schematic of MIE Locus. The distal promoter (dP), the major immediate early promoter (MIEP), intronic promoter 1 (iP1), and intronic promoter 2 (iP2) give rise to transcripts encoding full length IE1 and IE2. Transcription start sites for each promoter are depicted with raised arrows. **B)** Schematic of the 5’ ends up to and including exon 2 of IE transcripts derived from the MIE promoters. The translation start site (AUG) is marked in exon 2. Mature mRNAs encoding IE1 and IE2 will also include exons 3 and 4 or 3 and 5, respectively. Primer pairs designed to detect discreet transcripts by RT-qPCR (dP/MIEP = orange, iP1 = blue; iP2 = teal) are shown. A common reverse primer (black arrow) was used to amplify dP/MIEP and iP2-derived transcripts. **C)** THP-1 cells were infected with TB40/E WT HCMV (MOI = 2) and cultured for 5 days to promote the establishment of latency. At day 5, cells were treated with TPA (to promote macrophage differentiation and viral reactivation) or with a DMSO control. MIEP/dP, iP1-, and iP2-derived transcript accumulation was quantified relative to the low copy H6PD housekeeping gene by RT-qPCR. Data from three independent replicates is shown; error bars represent standard deviation.

**Fig 3.**
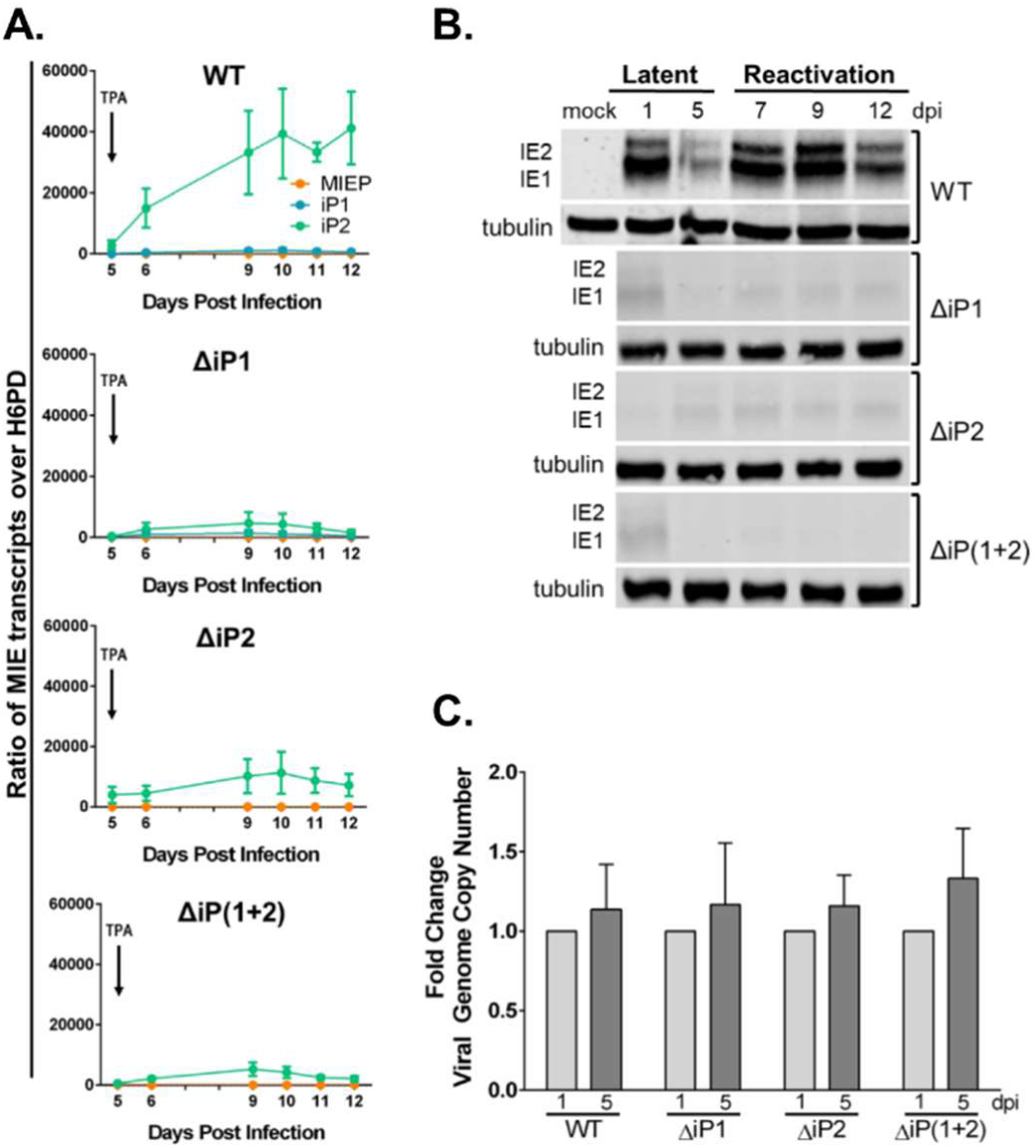
iP mutant viruses fail to re-express IE1 and IE2 following reactivation in THP-1 cells. THP-1 cells were infected with TB40/E WT, ΔiP1, ΔiP2 or ΔiP(1+2) HCMV (MOI = 2) and cultured for 5 days to promote the establishment of latency. At day 5, cells were treated with TPA or DMSO as a control to promote monocyte-to-macrophage differentiation and viral reactivation. **A)** RNA was isolated, and RT-qPCR was performed to monitor accumulation of iP1-, iP2-, and MIEP-derived transcripts relative to the low copy housekeeping gene H6PD. Data from three independent replicates is shown; standard deviation is depicted by error bars. **B)** Accumulation of IE1 and IE2 protein was measured during latency and following reactivation by immunoblotting using mouse monoclonal antibodies recognizing both IE1 and IE2. Tubulin was used as a loading control. A single experiment (representative of three independent experiments) is shown. **C)** Total DNA was isolated at days 1 and 5 during the latency period, and viral genomes were quantified by qPCR using a primer pair specific to the non-coding β2.7 region of the HCMV genome relative to BAC standard curve. Bars represent fold change over the number of viral genomes present at day 1 for each virus. Data from three independent replicates is shown; standard deviation is depicted by error bars.

To determine the significance of iP1 and iP2 for viral reactivation, we constructed recombinant viruses containing deletions surrounding the transcription start sites of iP1 (ΔiP1) and iP2 (ΔiP2) (Figure S3A). We also generated a mutant virus lacking both iP1 and iP2, ΔiP(1+2). Each mutant virus replicated with similar kinetics and to similar peak titers as the parental wildtype virus (Figure S3B), and efficiently expressed IE1 and IE2 (Figure S3C) during productive infection in fibroblasts. Together, these data indicate that the intronic MIE locus promoters are dispensable for expression of lytic cycle transactivators and for virus replication in fibroblasts.

We next compared the accumulation of transcripts derived from the MIEP, iP1, or iP2 during reactivation in THP-1 cells infected with wildtype (WT) or with each mutant virus (Figure 3A). Both iP1-and iP2-derived transcripts were diminished following reactivation in ΔiP1 infection. iP2-derived transcripts were diminished following reactivation in ΔiP2 infection; however, we could not measure iP1 transcript levels since the ΔiP2 mutation deleted the primer binding site for iP1-derived transcripts. Deletion of both iP1 and iP2 similarly diminished levels of iP2-derived transcripts following reactivation. MIEP-derived transcripts were detected at very low levels, if at all, in all infections. Consistent with the defect in IE1 and IE2 mRNA levels, deletion of iP1 and/or iP2 resulted in a striking decrease in accumulation of IE proteins after reactivation relative to the WT infection (Figure 3B). The failure of the recombinant viruses to reactivate could not be attributed to a defect in maintenance of latent infection, as viral genomes were maintained from 1 to 5 dpi with each virus (Figure 3C). From these results, we conclude that ΔiP mutant viruses establish a latent infection but are unable to reinitiate the expression of viral lytic cycle transactivators in response to a canonical reactivation stimulus, TPA.

Primary CD34+ HPCs are considered the ‘gold standard’ model for HCMV latency and reactivation (22–26). To evaluate whether iP1 and iP2 are required for reactivation in CD34+ HPCs, pure populations of infected (GFP+) CD34+ HPCs were isolated by cell sorting and seeded in long-term cultures over a stromal cell support to allow the establishment of latency (27). After 10 days, half of the culture was seeded by limiting dilution onto fibroblasts in a cytokine-rich media to promote differentiation and reactivation. The other half of the culture was mechanically lysed to quantify the amount of infectious virus produced prior to the reactivation event. The wildtype virus established latency and reactivated when co-cultured with fibroblasts, represented by a 3-fold increase in the frequency of infectious centers formed (Figure 4A). However, each of the ΔiP-mutant viruses exhibited a significant defect in reactivation. These results establish the requirement for the iP1 and iP2 promoters for efficient HCMV reactivation.

**Fig 4.**
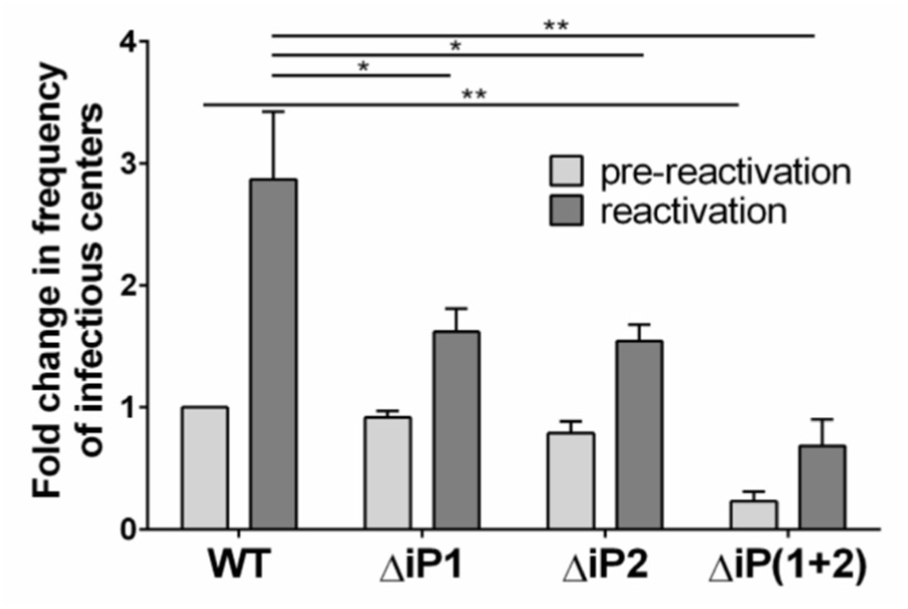
The intronic promoters are required for reactivation of HCMV from latency in CD34+ HPCs. CD34+ HPCs were infected with TB40/E WT, ΔiP1, ΔiP2, and ΔiP(1+2) expressing GFP as a marker for infection for 24 hours at an MOI of 2. Pure populations of infected (GFP+) CD34+ cells were isolated by FACS and maintained in long-term bone marrow culture for 10 days. Viable CD34+ HPCs were seeded by limiting dilution onto monolayers of permissive fibroblasts in a cytokine-rich media to promote myeloid differentiation (reactivation, dark gray). An equivalent number of cells was mechanically lysed and seeded in parallel to determine the infectious virus present in the culture prior to reactivation (pre-reactivation, light gray). The frequency of infectious centers formed pre-and post-reactivation was determined 14 days later by extreme limiting dilution analysis from the fraction of GFP+ wells at each dilution. Data is expressed as fold change over the frequency of infectious centers produced by the wildtype virus prior to reactivation. Data from three independent replicates is shown; standard error is depicted by error bars. Statistical significance was determined by multiple t tests comparing each mutant virus to the wildtype parental virus (* indicates a p value ≤ 0.05; ** indicates a p value ≤ 0.005).

Collectively, these findings indicate a significant shift away from the long-held paradigm that viral reactivation requires de-repression of the MIEP to drive IE1 and IE2 expression, and subsequent re-entry into the virus lytic cycle. Rather, we find that the majority of IE1 and IE2 expression upon reactivation is driven from recently identified promoters within intron A of the MIE transcriptional locus. Deletion of these promoters results in a striking defect in reactivation in both the THP-1 cell line and primary CD34+ HPC latency models. The cellular and/or viral factors that regulate intronic promoter activity during reactivation are currently unknown. Our data suggest that defining such factors will be critical to understanding the molecular events controlling HCMV latency and reactivation.

## Acknowledgments

The authors wish to acknowledge Dr. Jim Alwine and the laboratories of Dr. John Purdy and Dr. Anita Koshy at the University of Arizona for helpful discussion of the research presented in this manuscript. We acknowledge Mark Curry and the Arizona Cancer Center/Arizona Research Laboratories Division of Biotechnology Cytometry Core Facility for expertise and assistance with flow cytometry.

## Funding

Research reported in this publication was supported by the National Institute of Allergy and Infectious Diseases of the National Institutes of Health AI 079059 (F.G.), AI 143191 (F.G., N.M, J.K.), and a Pew Innovator Award (F.G). D.C-M is supported by a Postdoctoral Fellowship (18POST33960140) from the American Heart Association. J.B. is supported by a Postdoctoral Fellowship (129842-PF-16-212-01-TBE) from the American Cancer Society.

## Author contributions

Conceptualization, D.C-M., M.R., J.P.K., N.M., F.G.; Methodology, D.C-M., M.R., N.M., F.G.; Investigation, D.C-M., M.R., J.B., S.I-H.; Resources, N.M.; Writing – Original Draft, D.C-M and F.G.; Writing – Review & Editing, D.C-M., M.R., J.B., J.P.K., N.M. and F.G.; Visualization, D.C-M., N.M, F.G.; Funding Acquisition, J.P.K, N.M., and F.G.

## Competing interests

The authors declare no competing interests.

## Data and materials availability

All data is available in the main text or the supplementary materials.

## Supplementary Materials

Materials and Methods

Figures S1 to S3

Table S1

## References

1. Nogalski MT, Collins-McMillen D, Yurochko AD. 2014. Overview of Human Cytomegalovirus Pathogenesis., p. 15–28. In Yurochko AD, Miller WE (ed.), Human Cytomegaloviruses: Methods & Protocols. Humana Press New York.

2. Mocarski Jr E, Shenk T, Griffiths PD, Pass R. 2013. Cytomegaloviruses, p. 1960–2014. In Knipe DM, Howley PM (ed.), Fields Virology, 6th ed, vol. 2. Lippincott Williams & Wilkins, Philadelphia, PA, USA.

3. Stinski MF, Meier JL. 2007. Immediate-early viral gene regulation and function. In Arvin A, Campadelli-Fiume G, Mocarski E, Moore PS, Roizman B, Whitley R, Yamanishi K (ed.), Human Herpesviruses: Biology, Therapy, and Immunoprophylaxis. Cambridge University Press Copyright (c) Cambridge University Press 2007., Cambridge.

4. Goodrum F. 2016. Human Cytomegalovirus Latency: Approaching the Gordian Knot. Annual review of virology 3:333–357.

5. Collins-McMillen D, Buehler J. 2018. Molecular Determinants and the Regulation of Human Cytomegalovirus Latency and Reactivation. 10.

6. Thomsen DR, Stenberg RM, Goins WF, Stinski MF. 1984. Promoter-regulatory region of the major immediate early gene of human cytomegalovirus. Proceedings of the National Academy of Sciences of the United States of America 81:659–663.

7. Boshart M, Weber F, Jahn G, Dorsch-Hasler K, Fleckenstein B, Schaffner W. 1985. A very strong enhancer is located upstream of an immediate early gene of human cytomegalovirus. Cell 41:521–530.

8. Ghazal P, Lubon H, Fleckenstein B, Hennighausen L. 1987. Binding of transcription factors and creation of a large nucleoprotein complex on the human cytomegalovirus enhancer. Proceedings of the National Academy of Sciences of the United States of America 84:3658–3662.

9. Stinski MF, Roehr TJ. 1985. Activation of the major immediate early gene of human cytomegalovirus by cis-acting elements in the promoter-regulatory sequence and by virus-specific trans-acting components. Journal of virology 55:431–441.

10. Shelbourn SL, Kothari SK, Sissons JG, Sinclair JH. 1989. Repression of human cytomegalovirus gene expression associated with a novel immediate early regulatory region binding factor. Nucleic acids research 17:9165–9171.

11. Sinclair JH, Baillie J, Bryant LA, Taylor-Wiedeman JA, Sissons JG. 1992. Repression of human cytomegalovirus major immediate early gene expression in a monocytic cell line. The Journal of general virology 73 (Pt 2):433–435.

12. Stinski MF. 1999. Cytomegalovirus Promoter for Expression in Mammalian Cells, p. 211–233. In Fernandez JM, Hoeffler JP (ed.), Gene Expression Systems: Using Nature for the Art of Expression, 1st ed. Elsevier-Academic Press, Cambridge, Massachusetts.

13. Meier JL. 2001. Reactivation of the human cytomegalovirus major immediate-early regulatory region and viral replication in embryonal NTera2 cells: role of trichostatin A, retinoic acid, and deletion of the 21-base-pair repeats and modulator. Journal of virology 75:1581–1593.

14. Prosch S, Wuttke R, Kruger DH, Volk HD. 2002. NF-kappaB--a potential therapeutic target for inhibition of human cytomegalovirus (re)activation? Biological chemistry 383:1601–1609.

15. Keller MJ, Wu AW, Andrews JI, McGonagill PW, Tibesar EE, Meier JL. 2007. Reversal of human cytomegalovirus major immediate-early enhancer/promoter silencing in quiescently infected cells via the cyclic AMP signaling pathway. Journal of virology 81:6669–6681.

16. Yuan J, Liu X, Wu AW, McGonagill PW, Keller MJ, Galle CS, Meier JL. 2009. Breaking human cytomegalovirus major immediate-early gene silence by vasoactive intestinal peptide stimulation of the protein kinase A-CREB-TORC2 signaling cascade in human pluripotent embryonal NTera2 cells. Journal of virology 83:6391–6403.

17. Liu X, Yuan J, Wu AW, McGonagill PW, Galle CS, Meier JL. 2010. Phorbol ester-induced human cytomegalovirus major immediate-early (MIE) enhancer activation through PKC-delta, CREB, and NF-kappaB desilences MIE gene expression in quiescently infected human pluripotent NTera2 cells. Journal of virology 84:8495– 8508.

18. Arcangeletti MC, Vasile Simone R, Rodighiero I, De Conto F, Medici MC, Maccari C, Chezzi C, Calderaro A. 2016. Human cytomegalovirus reactivation from latency: validation of a “switch” model in vitro. Virology journal 13:179.

19. Lee CH, Lee GC, Chan YJ, Chiou CJ, Ahn JH, Hayward GS. 1999. Factors affecting human cytomegalovirus gene expression in human monocyte cell lines. Molecules and cells 9:37–44.

20. Yee LF, Lin PL, Stinski MF. 2007. Ectopic expression of HCMV IE72 and IE86 proteins is sufficient to induce early gene expression but not production of infectious virus in undifferentiated promonocytic THP-1 cells. Virology 363:174–188.

21. Arend KC, Ziehr B, Vincent HA, Moorman NJ. 2016. Multiple Transcripts Encode Full-Length Human Cytomegalovirus IE1 and IE2 Proteins during Lytic Infection. Journal of virology 90:8855–8865.

22. Goodrum FD, Jordan CT, High K, Shenk T. 2002. Human cytomegalovirus gene expression during infection of primary hematopoietic progenitor cells: a model for latency. Proc Natl Acad Sci USA. 99:16255–16260.

23. Petrucelli A, Rak M, Grainger L, Goodrum F. 2009. Characterization of a novel Golgi apparatus-localized latency determinant encoded by human cytomegalovirus. Journal of virology 83:5615–5629.

24. Umashankar M, Rak M, Bughio F, Zagallo P, Caviness K, Goodrum FD. 2014. Antagonistic determinants controlling replicative and latent states of human cytomegalovirus infection. J Virol. 88:5987– 6002.

25. Caviness K, Bughio F, Crawford LB, Streblow DN, Nelson JA, Caposio P, Goodrum F. 2016. Complex Interplay of the UL136 Isoforms Balances Cytomegalovirus Replication and Latency. mBio 7:e01986.

26. Buehler J, Zeltzer S, Reitsma J, Petrucelli A, Umashankar M, Rak M, Zagallo P, Schroeder J, Terhune S, Goodrum F. 2016. Opposing Regulation of the EGF Receptor: A Molecular Switch Controlling Cytomegalovirus Latency and Replication. PLOS pathogens 12:e1005655.

27. Umashankar M, Goodrum F. 2014. Hematopoietic long-term culture (hLTC) for human cytomegalovirus latency and reactivation. Methods in molecular biology 1119:99–112.

