## Supplementary material for "Alternative Promoters Drive Human Cytomegalovirus Reactivation from Latency"

### Supplementary Materials

#### Materials and Methods

**Cells**. The human monocytic leukemia cell line, THP-1, was purchased from ATCC (Manassas, VA). THP-1 cells were cultured in Roswell Park Memorial Institute (RPMI) 1640 medium supplemented with 10 % fetal bovine serum (FBS), 2 mM L-alanyl-glutamine, 2 mM sodium pyruvate, 0.05 mM β-mercaptoethanol, 100 U/mL penicillin, and 100 μg/mL streptomycin. Cells were maintained below 1 million cells per mL. Human primary embryonic lung fibroblasts (MRC-5; purchased from ATCC) were cultured in Dulbecco's modified Eagle's medium (DMEM) supplemented with 10% FBS, 10 mM HEPES, 2 mM L-alanyl-glutamine, 1 mM sodium pyruvate, 0.1 mM non-essential amino acids, 100 U/mL penicillin, and 100 μg/mL streptomycin. CD34+ human progenitor cells (HPCs) were obtained from medical waste from normal bone marrow harvests at the University of Arizona Medical Center. All waste containers were de-identified prior to receipt of specimens. CD34+ HPCs were isolated using a CD34 MicroBead kit according to manufacturer's instructions (magnetically activated cell sorting or MACS; Miltenyi Biotech, San Diego, CA). Pure populations of CD34+ HPCs were cultured in Myelocult H5100 (Stem Cell Technologies, Cambridge, MA) supplemented with hydrocortisone, 100 U/mL penicillin, and 100 μg/mL streptomycin and maintained in long-term co-culture with M2-10B4 and S.l./S.l. murine stromal cell lines (Stem Cell Technologies).

**Viruses**. The HCMV TB40/E bacterial artificial chromosome (BAC) was previously engineered to express green fluorescent protein (GFP) as a visual marker for infection (1, 2). The ΔiP1, ΔiP2, and ΔiP(1+2) mutant bacterial artificial chromosome (BAC) clones were made by deletion of large fragments in MIE intron A surrounding the transcription start sites of each of the iP-derived transcripts. A cassette of *galK* flanked by *loxP* sites was inserted into the TB40/E wildtype parental BAC (substituting for the sequence surrounding each promoter) via homologous recombination in SW102 *Escherichia coli*. BACs underwent a round of positive selection for *galK*, after which *galK* was floxed out, leaving the deletions. 323 bp (206968 – 207291) were deleted from ΔiP1, 410 bp (206567 -206977) from ΔiP2, and 733 bp (206567 – 207291) from ΔiP(1+2). The desired mutations were confirmed by sequencing. Infectious virus (P0) was produced by transfecting BAC genomes (20 μg) and 2 μg of an expression plasmid encoding UL82 (pp71) into 5 million MRC-5 fibroblasts, which were maintained until 100% cytopathic effect (CPE) was observed. Virus stocks (P1) were produced by infecting MRC-5 fibroblasts with the P0 stock, then purifying cell-free virus by density gradient centrifugation through a 20% D-sorbitol cushion when cells reached 100% CPE. P1 stocks were resuspended in Iscove's modified DMEM supplemented with 2% bovine serum albumin (BSA) and stored at -80°C. Infectious virus yields were determined by 50% tissue culture infective dose ($TCID_{50}$) on MRC-5 fibroblasts (3).

**THP-1 latency model**. THP-1 cells were infected as monocyte-like suspension cells at a density of 0.5 million cells per mL and an MOI of 2 (as determined by $TCID_{50}$ using MRC-5 fibroblasts). To maximize infection efficiency, infected cell suspension was mixed by rocking every 30 minutes for 4 hours, followed immediately by a "spinfection" at 450 x g for 20 minutes. Cells were cultured for 5 days post infection in non-tissue culture-treated six well plates (1.5

million cells in 3 mL per well). On day 5, cells from each experimental group were pooled and centrifuged at 120 x g for 7 minutes, then resuspended at 0.5 million cells per mL. Cells were treated with 100 µM 12-O-Tetradecanoylphorbol-13-acetate (TPA) and plated on tissue culture-treated plates to promote monocyte-to-macrophage differentiation (and viral reactivation) or treated with DMSO as a solvent control and cultured in non-tissue culture-treated six well plates (1.5 million cells in 3 mL per well).

**Immunoblotting**. Whole cell lysates were collected in RIPA buffer (25mM Tris-HCl pH 6.7, 150mM NaCl, 1% octylphenoxypolyethoxyethanol, 1% Na deoxycholate, 0.1% sodium dodecyl sulfate) supplemented with 1x Halt Protease Inhibitor (Thermo Scientific, Waltham, MA) and 2 mM phenylmethylsulfonyl fluoride (PMSF) for 30 minutes on ice. Lysates were stored at -20°C until analyzed. Total protein concentration for each sample was determined by BCA assay (Thermo Scientific), and samples were denatured by boiling in protein lysis buffer (120 mM Tris pH 6.8, 4% glycerol, 100mM dithiothreitol, 0.01% Bromophenol blue) for 10 minutes. A total of 50 µg of protein per sample was loaded for each lane. Proteins were separated by SDS-PAGE and transferred to polyvinylidene difluoride (PVDF) membranes. Blots were blocked in Odyssey Blocking Buffer (Tris Buffered Saline) (LI-COR, Lincoln, NB) for 45 minutes, then probed with monoclonal antibodies against IE1/2 (generous gift from Tom Shenk) and tubulin (Sigma-Aldrich, St. Louis, MO) overnight at 4°C. Blots were rinsed with TBS-Tween (1x Tris Buffered Saline, 20% Tween 20, 1 g/L bovine serum albumin, 0.02% sodium azide) then probed with fluorescently labeled secondary antibodies (LI-COR) for 1 hour at room temperature. Proteins were detected using an Odyssey infrared imaging system (LI-COR).

**Reverse transcriptase quantitative polymerase chain reaction (RT-qPCR)**. Total RNA was extracted using a Zymo Duet RNA/DNA isolation kit (Zymo Research, Irvine, CA). cDNA was synthesized using the Transcriptor First Strand cDNA Synthesis Kit (Roche, Basel Switzerland). Briefly, total RNA (400ng) was combined with 2.5 µM Anchored-oligo$(dT)_{18}$ Primers and denatured at 65°C for 10 minutes. A Reverse Transcriptase (RT) master mix (1x Transcriptor RT Reaction Buffer, 40 U/µL Protector RNase Inhibitor, 10 mM Deoxynucleotide Mix, 20 U/µL Transcriptor RT) was added to the template-primer mix (a no RT- control was made by substituting water for RT in a single reaction). Samples were incubated in a Mastercycler (Eppendorf, Hamburg, Germany) for 60 minutes at 50°C, then raised to 85°C for 5 minutes to inactivate the Reverse Transcriptase. Final reaction products were diluted 1:4 in PCR water to reduce the concentration of $MgCl_2$. The resulting single-stranded cDNA was amplified in a quantitative polymerase chain reaction (qPCR) using 1x Lightcycler 480 SYBR Green I Master Mix (Roche) and 0.5 µM of a series of sequence-specific primer pairs (see Table S1 for detailed target sequences). Relative expression of each mRNA was calculated using the Pfaffl method that accounts for efficiency of individual primer pairs (4). Primer efficiencies were calculated using an internal standard curve made with cDNA from lytically infected fibroblasts.

**Multistep virus growth curves**. Viral growth kinetics were measured by two-step low MOI growth curve. MRC-5 cells were infected at an MOI of 0.02 and cells and medium were collected over a 16 day time course of infection. Virus titers were determined for each time point by quantification of $TCID_{50}$ in MRC-5 fibroblasts (2).

**Quantitative PCR for measuring viral genomes**. Total DNA was isolated at the indicated time points using a Zymo Duet RNA/DNA isolation kit (Zymo Research). Viral genomes were quantified by quantitative polymerase chain reaction (qPCR) using 1x Lightcycler 480 SYBR Green I Master Mix (Roche) and primers targeted against the genomic region corresponding to the non-coding β2.7 RNA. Number of viral genomes present in each sample was quantified relative to BAC standard curve.

**Assays of infectious centers for latency**. Frequency of reactivation for each mutant virus relative to wildtype in CD34+ human progenitor cells (HPCs) was quantified as previously described (5, 6). Briefly, pure populations of CD34+ human progenitor cells (HPCs) were infected with WT, ΔiP1, ΔiP2, or ΔiP(1+2) virus (MOI = 2) expressing GFP as a marker for infection. At 24 hpi, infected (GFP+) CD34+ cells were isolated by fluorescence activated cell sorting (FACS) and incubated in long-term culture with a stromal cell support to maintain latency. At 10 dpi, viable CD34+ HPCs were seeded by limiting dilution onto monolayers of permissive MRC-5 fibroblasts (reactivation). An equivalent number of CD34+ HPCs were mechanically disrupted and seeded in parallel to quantify infectious virus present prior to reactivation (pre-reactivation). Frequency of infectious centers formed prior to and following reactivation was quantified 14 days later by extreme limiting dilution analysis (ELDA) of GFP+ wells.


1. **Sinzger C, Hahn G, Digel M, Katona R, Sampaio KL, Messerle M, Hengel H, Koszinowski U, Brune W, Adler B.** 2008. Cloning and sequencing of a highly productive, endotheliotropic virus strain derived from human cytomegalovirus TB40/E. The Journal of general virology **89:**359-368.
2. **Umashankar M, Petrucelli A, Cicchini L, Caposio P, Kreklywich CN, Rak M, Bughio F, Goldman DC, Hamlin KL, Nelson JA, Fleming WH, Streblow DN, Goodrum F.** 2011. A novel human cytomegalovirus locus modulates cell type-specific outcomes of infection. PLoS pathogens **7:**e1002444.
3. **Petrucelli A, Rak M, Grainger L, Goodrum F.** 2009. Characterization of a novel Golgi apparatus-localized latency determinant encoded by human cytomegalovirus. Journal of virology **83:**5615-5629.
4. **Pfaffl MW.** 2001. A new mathematical model for relative quantification in real-time RT-PCR. Nucleic acids research **29:**e45.
5. **Goodrum FD, Jordan CT, High K, Shenk T.** 2002. Human cytomegalovirus gene expression during infection of primary hematopoietic progenitor cells: a model for latency. Proc Natl Acad Sci USA. **99:**16255-16260.
6. **Umashankar M, Goodrum F.** 2014. Hematopoietic long-term culture (hLTC) for human cytomegalovirus latency and reactivation. Methods in molecular biology **1119:**99-112.

**Figure S1.**

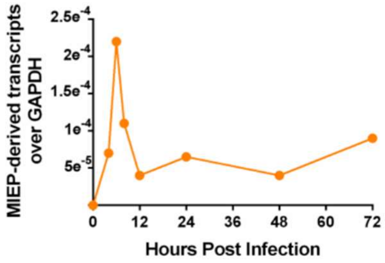


**Amplification of MIEP-derived transcripts in permissive fibroblasts.** MRC-5 fibroblasts were infected with WT TB40E HCMV (MOI = 1) and cultured for three days post infection. Total RNA was isolated at the indicated time points and RT-qPCR was performed to monitor the accumulation of MIEP-derived transcripts. Transcripts are expressed as a ratio over the high copy housekeeping gene GAPDH. A single replicate (representative of three independent experiments) is shown.

**Figure S2.**

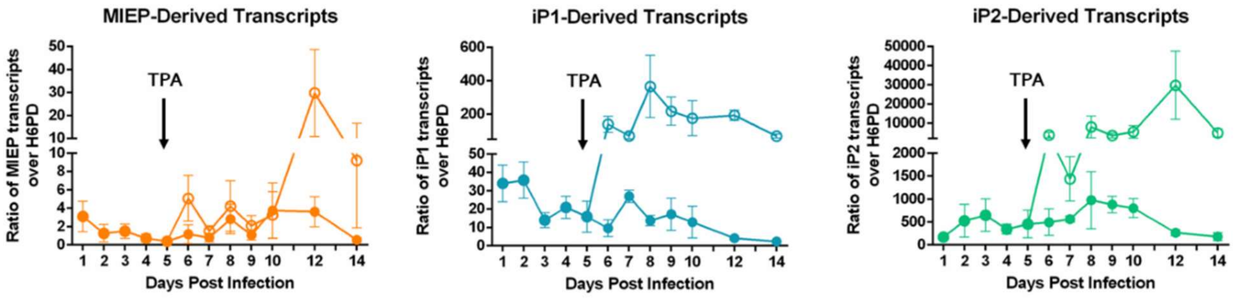


**MIEP-derived and iP-derived transcripts on their individual scales during reactivation.** THP-1 cells were infected with WT HCMV (MOI = 2) and cultured for 5 days to promote the establishment of latency. At day 5, cells were treated with TPA (to promote macrophage differentiation and viral reactivation) or with a DMSO control. RNA was isolated, and RT-qPCR performed to monitor accumulation of MIEP-derived (left panel), iP1-derived (center panel), and iP2-derived (right panel) transcripts. Transcripts are expressed as a ratio over the low copy housekeeping gene H6PD. Data from three independent replicates is shown; standard deviation is depicted by error bars.

**Figure S3.**

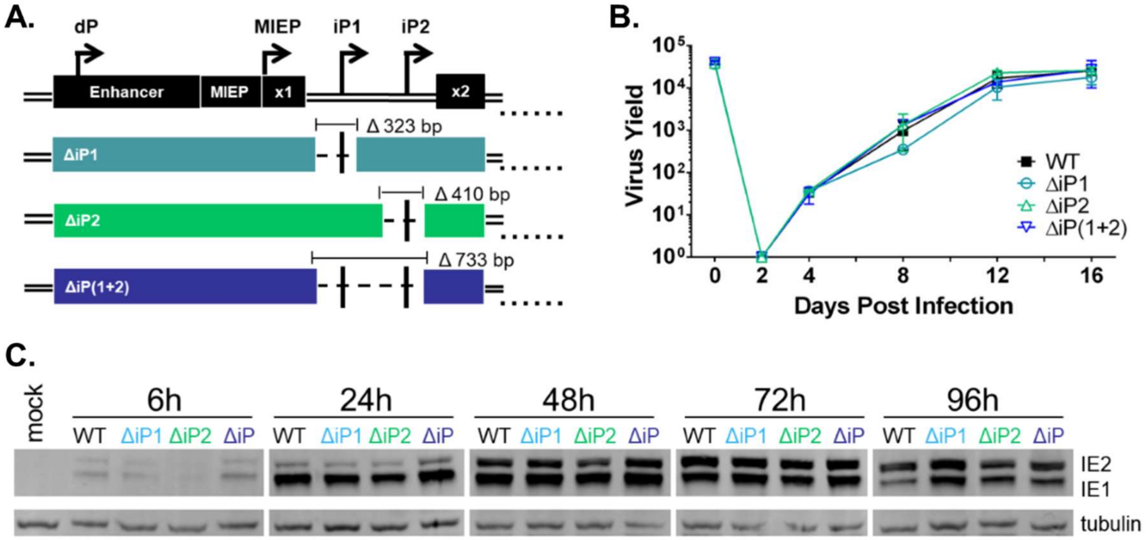


**Construction & characterization of iP mutants. A)** HCMV ΔiP mutants were made using BAC-mediated recombineering. Inserts (*galK* flanked by *loxP* sites) were inserted into the TB40E wildtype parental BAC via homologous recombination at the various promoter sites. BACs underwent a round of positive selection for *galK*, after which *galK* was floxed out resulting in deletions of 323 bp (ΔiP1), 410 bp (ΔiP2), or 733 bp (ΔiP(1+2)) surrounding the transcription start sites of the iP transcripts. **B)** Two-step low MOI dose curves were performed. MRC-5 fibroblasts were infected with WT, ΔiP1, ΔiP2 or ΔiP(1+2) HCMV (MOI = 0.02). Cell lysates were collected at the indicated days post infection, and TCID50 was performed to quantify viral replication and construct growth curves. Data from three independent replicates is shown; standard deviation is depicted by error bars. **C)** MRC-5 fibroblasts were infected with WT, ΔiP1, ΔiP2, or ΔiP(1+2) HCMV. Whole cell lysates were collected, and Western blot analysis was performed to monitor IE1 and IE2 expression. Tubulin was used as a loading control. A single replicate (representative of three independent experiments) is shown.

**Table S1. qPCR primers.**

| Transcript | Primer Pair |
|---|---|
| MIEP/dP-derived | FWD: 5′-TTG ACC TCC ATA GAA GAC AC-3″ |
| | REV: 5′-AGG ACT CCA TCG TGT CAA GG-3′ |
| iP1-derived | FWD: 5′-CTT AAG GCA GCG GCA GAA-3′ |
| | REV: 5′-CAA GGA CGG TGA CTG ACT C-3′ |
| iP2-derived | FWD: 5′-TAG CTG ACA GAC TAA CAG AC-3′ |
| | REV: 5′-AGG ACT CCA TCG TGT CAA GG-3′ |
| IE1 (UL123) | FWD: 5′-TGA CCG AGG ATT GCA ACG A-3′ |
| | REV: 5′-CCT TGA TTC TAT GCC GCA CC-3′ |
| IE2 (UL122) | FWD: 5′-CAG AAC TCG GGT GAC ATC CT-3′ |
| | REV: 5′-CCG GTC CTA CTG GAA TCG-3′ |
| H6PD | FWD: 5′-GGA CCA TTA CTT AGG CAA GCA-3′ |
| | REV: 5′-CAG GGT CTC TTT CAT GAT GAT CT-3′ |
| **DNA** | **Primer Pair** |
| β2.7 | FWD: 5′-TGT TCT TCT GGT TCA TTT CCT ATG-3′ |
| | REV: 5′-CGT GTC CGG TCC TGA TTC-3′ |
